# Digitally Deconstructing Leaves in 3D Using X-ray Microcomputed Tomography and Machine Learning

**DOI:** 10.1101/814954

**Authors:** Guillaume Théroux-Rancourt, Matthew R. Jenkins, Craig R. Brodersen, Andrew McElrone, Elisabeth J. Forrestel, J. Mason Earles

## Abstract

**Premise of the study:** X-ray microcomputed tomography (microCT) can be used to measure 3D leaf internal anatomy, providing a holistic view of tissue organisation. Previously, the substantial time needed for segmenting multiple tissues limited this technique to small datasets, restricting its utility for phenotyping experiments and limiting our confidence in the conclusion of these studies due to low replication numbers.

**Methods and Results:** We present a Python codebase for random-forest machine learning segmentation and 3D leaf anatomical trait quantification which dramatically reduces the time required to process single leaf microCT scans into detailed segmentations. By training the model on each scan using 6 hand segmented image slices out of >1500 in the full leaf scan, it achieves >90% accuracy in background and tissue segmentation.

**Conclusion:** Overall, this 3D segmentation and quantification pipeline can reduce one of the major barriers to using microCT imaging in high-throughput plant phenotyping.

## INTRODUCTION

Leaves are complex and highly sophisticated 3D geometries optimized over the course of evolutionary time to balance water distribution, photosynthesis, and structural integrity, among many other biological functions. Yet only recently has imaging technology enabled a clear view and, more importantly, the capacity to digitally represent leaf 3D anatomy (Théroux-Rancourt et al., 2017). Today, 3D imaging permits precise spatial measurement and biophysical modeling of leaf internal geometry that can deliver novel insights about basic leaf function, such as CO_2_ transport (Ho et al., 2016; Lehmeier et al., 2017; Earles et al., 2018, 2019; Lundgren et al., 2019), H_2_O transport (Scoffoni et al., 2017), and mechanical structure (Pierantoni et al., 2019). Embracing the 3D complexity of leaf geometry permits us to understand when dimensionality reduction is tolerable and will ultimately guide more precise mechanistic scaling from tissue to crop/ecosystem.

Computationally, 3D imaging often produces large datasets (>20 Gb) with hundreds to thousands of digital cross sections that do not immediately yield biologically relevant information. Regardless of the imaging modality, 3D images must be subsequently processed to extract biologically relevant information, such as tissue type, chemical composition, and material type. In the case of X-ray micro-computed tomography (microCT) applied to plant leaves, this has led to the 3D description of the complex organization of the mesophyll cells and their surface area (Ho et al., 2016; Théroux-Rancourt et al., 2017), and the description of novel anatomical traits related to the intercellular airspace (Lehmeier et al., 2017; Earles et al., 2018). Tissue segmentation can be done quickly using both proprietary and open source software via 3D thresholding based on pixel intensity values. However, in the case of leaf microCT scans, pixel intensity can primarily, and most often solely, distinguish between water-filled cells and air-filled void areas. As such, quick segmentations can generally only label cells and airspace, especially when using phase contrast reconstruction (Théroux-Rancourt et al., 2017), resulting in the different tissues of a leaf, e.g. the epidermis, the mesophyll cells, the bundle sheaths, and the veins being grouped together. Using this method, there is not a clear distinction between the background and the intercellular airspace. This results in segmentations being limited to small leaf volumes consisting solely of mesophyll cells and airspace to estimate leaf porosity and cell surface area, traits commonly measured when related to photosynthetic efficiency (e.g. Ho et al., 2016). However, small leaf volumes do not necessarily represent the whole leaf trait average, and as such a larger volume including veins is needed to limit sampling bias. To separate the leaf from the background and segment the different tissues within the leaf, current applications generally rely on the onerous process of hand-segmentation, i.e. drawing with a mouse or a graphic tablet over single slices of a microCT scan to delimit and assign a unique value to each of the different tissues, either slice-by-slice or through the interpolation between different delimited regions throughout the scan (Théroux-Rancourt et al., 2017; Harwood et al., 2020). As a result, studies incorporating 3D microCT datasets have been limited to smaller scanning endeavors, and the low replicability of these studies limits the impact of conclusions therein. Hand segmentation, as described above, can take up to one day of work for a coarse scale segmentation of tissues other than mesophyll cells and airspace (as in Théroux-Rancourt et al., 2017). Further, highlighting natural variations in size and curvature of the various tissues can substantially increase hand segmentation time (see for example Harwood et al. (2020) on a similar issue using serial block face scanning electron microscopy). Hence, segmentation is currently a major bottleneck in the use of this technology.

Machine learning (ML) presents an opportunity to substantially accelerate the image segmentation process for plant biological applications. Conventional computer vision techniques rely on a human to engineer and select visual features, such as shape, pixel intensity, and texture, that ultimately guide the underlying segmentation process. On the other hand, ML-based image processing allows the machine to directly select or engineer visual features (e.g. Çiçek et al. 2016, Berg et al. 2019). ML-based image processing techniques fall along a continuum of unsupervised to supervised learning, which defines the degree to which the machine uses ground-truth data for guiding its optimization function. Given the large number of images generated during an X-ray microCT scan, ML-based image processing could lead to major efficiency gains in terms of human effort, enabling higher sample throughput and more complete data utilization as outlined above. In this study, we present a random forest ML framework for image segmentation of single X-ray microCT plant leaf scans and test its performance on an a grapevine leaf scan (for an in depth user manual, please refer to the repository of this program: github.com/plant-microct-tools/leaf-traits-microct). We finally demonstrate how the rich 3D output can be used to extract biologically meaningful metrics from these segmented images.

## METHODS AND RESULTS

### Random forest segmentation and leaf traits analysis pipeline

The following pipeline was built for our projects using X-ray synchrotron-based microCT imaging and uses freely available and open source software ImageJ (Schneider et al., 2012) and the Python programming language for machine-learning segmentation and for image analysis. Synchrotron-based imaging allows to reconstruct the scans using the gridrec reconstruction, which reflects X-ray absorption and provides a sharp but low contrast image highlighting the interface between cells (Dowd et al., 1999), and the phase contrast reconstruction, providing images with increased contrast between material of different absorptance (Paganin et al., 2002). Both reconstructions were at the base of our previous method (Théroux-Rancourt et al., 2017). In its current state, the program needs the gridrec and phase contrast reconstructions in 8-bit depth (Figure 1). To prepare for model training and automated segmentation, one needs to prepare hand labelled slices. Briefly, using ImageJ, we first binarize, i.e. convert to black and white, the two reconstructions by applying a threshold, where grayscale values below are considered air and above are considered cells (Figure 1). Those two binary stacks are combined together as in Théroux-Rancourt et al. (2017). Hand labelling is then done directly in ImageJ by drawing around each tissue, repeating the labelling over the desired number of slices while paying attention to cover a range of anatomical variations such as the density and orientation of veins, the most important cause of variation between slices (Théroux-Rancourt et al., 2017). In the current case the background, both epidermises, and the bundle sheaths and veins pairs (1-3 per slice, each tissue labeled separately) were labelled by a single individual, and consistency was validated by the first author (Figure 1; see also Supplementary Table S1 for the number of pixels per class). For a detailed methodology on preparing hand labelled slices, please refer to the repository of this program (github.com/plant-microct-tools/leaf-traits-microct).

**Figure 1.**
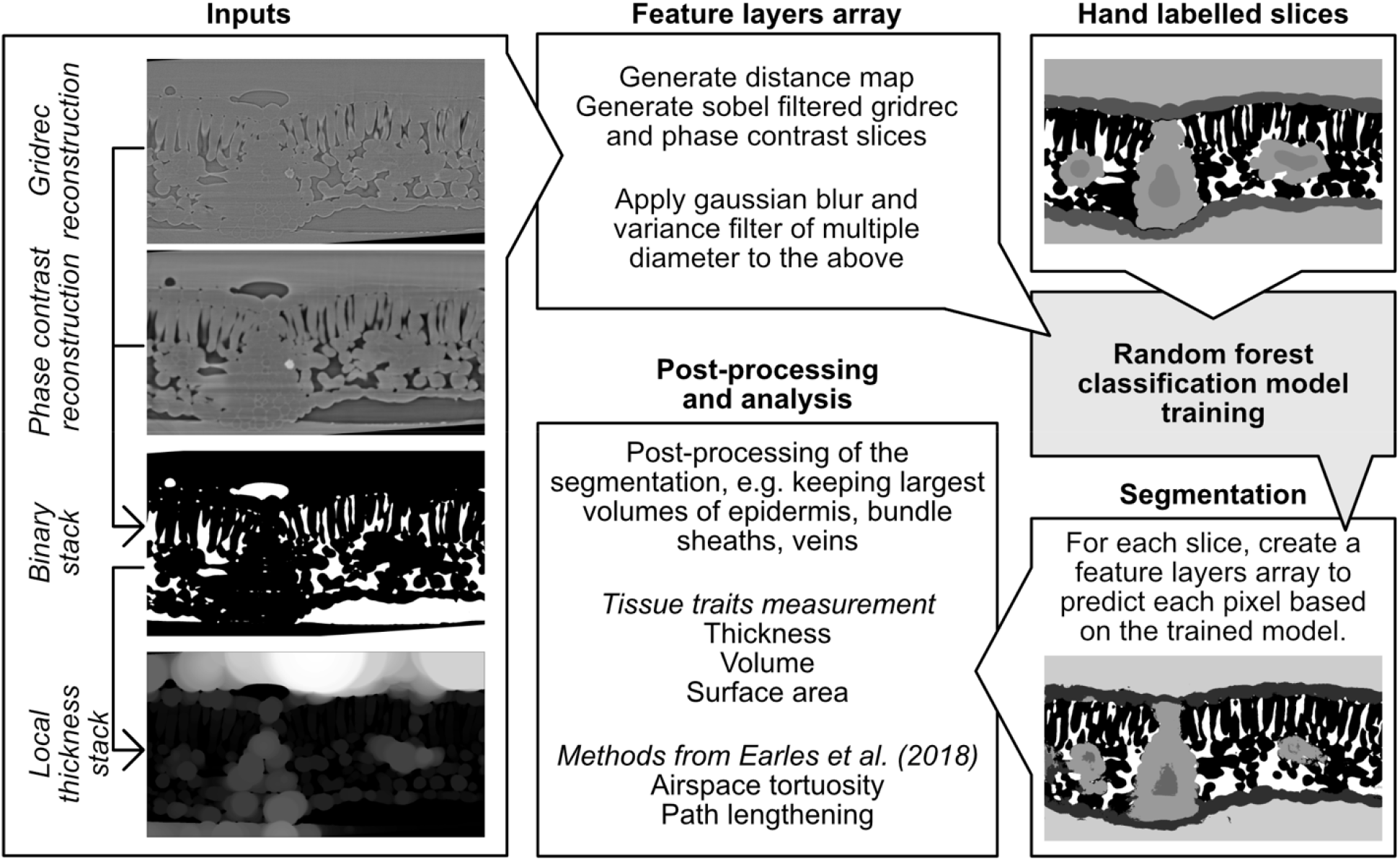
Schematic of the segmentation and analysis pipeline. Reconstructed microCT scans are manually thresholded to find the best value to segment the airspace of the leaf (as in Théroux-Rancourt et al. 2017). Using this binary stack, a local thickness stack is created, which identifies for each pixel the diameter of the largest sphere contained in that area (lighter pixel values mean larger diameters). These inputs stacks are used to generate the feature layers arrays needed, along with the hand labelled slices, for the random forest classification model training. With the trained model, the complete stack of images is predicted, and from this predicted stack the image is post-processed to remove false classifications, and leaf traits are analyzed. Note that all images are from the same position within the stack (i.e. same slice) except for the segmented image: the same slice as was hand labelled provides identical segmentation.

The random-forest classification model can then be trained using the hand labelled slices. We used a custom Python 3.7 program using numpy (Oliphant, 2006) for data structure, scikit-image (van der Walt et al., 2014) for image processing, and scikit-learn (Pedregosa et al., 2011) for random-forest machine learning functions. The image processing and random-forest classification is summarized in Figure 1. First, using the gridrec and phase contrast reconstructions the program creates a binary image (as defined above) using the threshold values for both stacks as input variables. This binary image is then used to create a local thickness map, which identifies through distance transformation and for each pixel the diameter of the largest sphere contained in that area, i.e. representing an estimate of the axis length or diameter of that structure or cell. This map provides additional information needed to predict class based on the object 3D size. Feature layer arrays are then built by applying a gaussian blur or a variance filter, both of different diameters, to the gridrec and phase contrast slices. The same filters are also applied to a map of the distance from the top and bottom edge of the image to its center, and to gridrec and phase contrast slices that have been sobel filtered to emphasize edges. These feature layer arrays are then used, along with the local thickness map, to train the random-forest classification model by predicting pixel values on the desired number of hand labelled training slices, which are randomly selected within the full hand labelled stack. A default of 50 estimators is used and can be adjusted by the user. Further parameters used to train the random forest are whether to use out-of-box samples to estimate accuracy (hard-coded to “True”) and whether to reuse the solution of the previous call to fit and add more estimators to the ensemble, or just fit a new forest (hard-coded to “False”, i.e. a new forest is always fitted).

After training the model and testing its accuracy on the hand labelled slices, it is used to automatically predict the remaining slices of the microCT scan, (generally > 1500) in a few hours, a massive improvement over conventional methods which require hundreds of hours of manual work (Théroux-Rancourt et al., 2017). In this article we present results for one single scan. If users intend to segment multiple scans, the procedure above, i.e. hand labeling and model training, would have to be done for each scan as the framework currently provides suitable and accurate segmentations for plant anatomy analysis when trained on each scan.

The full stack prediction can be then passed on to the leaf traits analysis pipeline. A first step is to identify all tissues and apply post-prediction correction to remove some false predictions that would bias biological trait analysis. In the case of laminar leaves, his includes identifying the two largest epidermis structures of similar volumes, as the abaxial and adaxial epidermis should each form a single volume within the stack. Thus smaller volumes labelled as epidermis, for example identified within the mesophyll cells, are not considered to be actual epidermis and are removed during this correction. For veins and bundle sheaths, as they are also usually highly connected to each other, small volumes do not represent actual tissues and as such very small and unique volumes, generally below 27 px^3^, are removed. Hence, the stack used for trait analysis was corrected to include biologically relevant volumes.

From this corrected stack, biological metrics are computed. Here, we measured the thickness at each point along the leaf surface for the whole leaf, the abaxial and adaxial epidermis, the whole mesophyll (leaf without the epidermis), all of which include standard deviation. We also measured the volume of all segmented tissues through a voxel count, as well as the surface area of the mesophyll cells connected to the airspace through a marching cube algorithm. Further analysis of the airspace can be made to compute tortuosity and path lengthening using a python-version of Earles et al. (2018) methods, which are not included in the current methods analysis.

### Testing the segmentation program

To test the performance of the segmentation program, we use the microCT scan of a ‘Cabernet sauvignon’ grapevine (*Vitis vinifera* L.) leaf acquired at the TOMCAT tomographic beamline of the Swiss Light Source (SLS) at the Paul Scherrer Institute in Villigen, Switzerland. Samples were prepared for microCT scanning as in Théroux-Rancourt et al. (2017), the sample was mounted between pieces of polyimide tape and fixed upright in a styrofoam holder, and 1801 projections of 100 ms were acquired at 21 keV over 180° total rotation using a 40x objective, for a final pixel size of 0.1625 μm. The scans were reconstructed with the gridrec and paganin algorithms using the reconstruction pipeline at the TOMCAT beamline. Twenty-four slices spread evenly across the full 1920 slices stack were hand labelled for epidermis, background, veins, and bundle sheaths, mesophyll cells, and intercellular airspace as briefly described above. To facilitate the testing, the *x* and *y* dimensions were halved, yielding a pixel size of 0.325 μm in those dimensions, but keeping the original dimensions in the depth (*z*) dimension, hence reducing the file size by four down to 1.5 Gb, a size easily handled by the program.

To understand the impact of training a model using different numbers of manually segmented slices, we iteratively trained the model using 1 through 12 slices. We repeated this process 25 times for each number of training slices using randomly selected training slices for each iteration. To cross-validate between hand-labeled ground truth and model predictions, each trained model was used to predict a test set consisting of all slices that were hand labelled but that had not been used to train the model (e.g. if six slices were used for training, the remaining 18 slices were predicted using the model). Based on the model generated from the training procedure, we performed segmentation on at least 5 entire stacks (see below). These fully segmented stacks were analyzed for their leaf level anatomical predictions. Confusion matrices were then created for each prediction test. Note that no post-prediction corrections were applied and as such the results below present the raw predictions.

From each confusion matrix, we evaluated precision and recall for each biological class (Figure 2). In the context of automated information retrieval for microCT image segmentation, recall may be interpreted as the sensitivity of the trained model to a given pixel class, i.e. the portion of correctly identified pixels in a given class, relative to all pixels belonging to this class. On the other hand, precision represents the positive predictive value of the model within a given pixel class, i.e. the number of pixels correctly identified as belonging to a given class, divided by this value plus the number of pixels falsely identified. It can be logically deduced why some people refer to recall as *quantity* of positive identification, and precision as the *quality* of positive identification.

**Figure 2.**
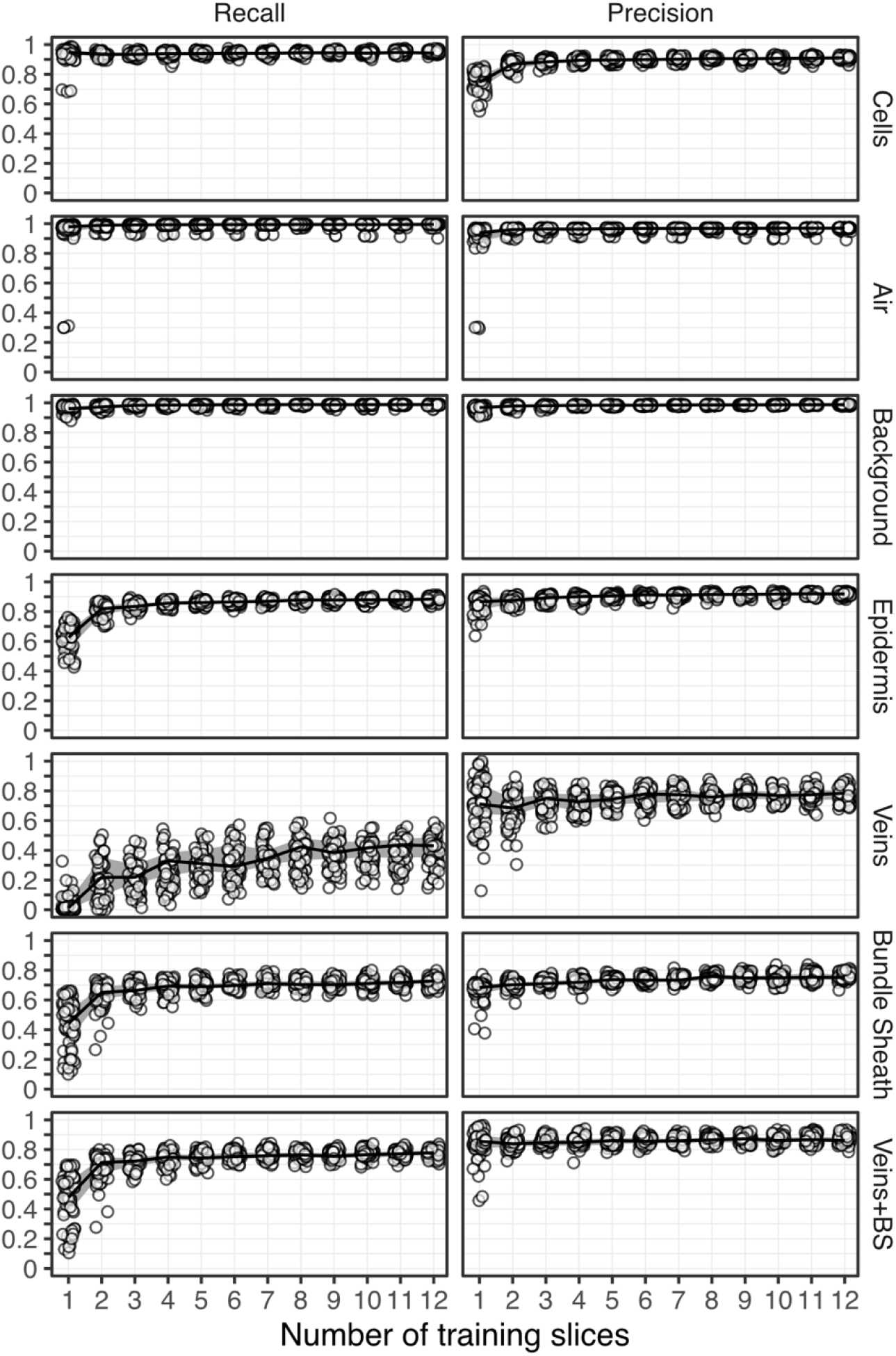
Recall (left) and precision (right) as a function of the number of training slices used to predict tissue classes in a microCT leaf scan (pixel size: 0.325 μm; 762 999 pixels per slice to predict). Solid lines represent the median value of 30 prediction per amount of training slices, and gray ribbons represents the 25th and 75th quantiles.

In the mesophyll cell class, recall was generally > 90% even when training on < 3 manually segmented slices, meaning that > 90% of all mesophyll cell pixels were correctly identified as cells, suggesting the trained random forest model is highly sensitive to cells. The same can be said for the airspace and background classes, which plateau at about 95% recall using >1 training slice. The trained models do not appear to be as sensitive to pixels of the epidermis class. Indeed, we observed a minimum of 4 training slices required to drive epidermis class recall above 90%. With vein and bundle sheath considered together as one class, at least 4 training slices were required to reach a maximum recall value of ~80%; the remaining 20% were false negatives, i.e. identified as other classes. Interestingly, when separating bundle sheath and vein into distinct classes, the bundle sheath class also reaches a maximum recall value of ~75% using >4 training slices. Isolating the vein class from bundle sheath, greatly impacts the trained model’s sensitivity to detection of vein. Recall was not observed above 55%, and generally stayed under 40% unless the model was trained on > 8 manually segmented slices.

To achieve precision > 90% in the airspace, background, and epidermis tissue classes a minimum of 2 training slices should be used. Interestingly, training on > 2 slices did not seem to translate to a substantial improvement in precision for these classes. However, observed precision for the mesophyll cell tissue class did not plateau until training on > 3 slices. While the maximum precision for mesophyll cells was stably > 90%, the lower precision values consistently observed when training on 1 or 2 slices, as low as 60%, suggest the software is not as reliable for this tissue class. So, it is important to train on > 3 slices if mesophyll cell traits are of importance. In the vein class, the software was observed to positively identify pixels at a rate of about 80% when trained on > 2 slices. In other words, even though the software is not very sensitive to the vein class, it is quite reliable when it does make a positive identification in the vein class.

To evaluate how the number of training slices affected the measurement of biological traits, we carried out at least five supplementary model training per number of training slices and made predictions over the full stack instead of over the 24 hand labelled slices. These full stack predictions were then passed through the leaf traits analysis program to extract relevant leaf anatomical traits (Figure 3). Anatomical measures were the least constant between predictions when using one training slice. The most variable were the epidermises thickness estimates, with values near 0 μm for the abaxial epidermis, or close to 30 μm in the adaxial epidermis, meaning that false segmentations of epidermis occurred between both epidermises such that they were connected and could not be automatically distinguished from one another as happened from 3 training slices onward. This false segmentation of the epidermis led to a highly variable whole mesophyll thickness (i.e. the leaf without the epidermis), which became less variable (< 5%) when using at least 3 training slices. However, the overall leaf thickness was the least variable, with less than ~1.5 μm variation (~1% total thickness) when using 3 or more training slices, a variation we consider equal or even lower than manual measures. This technique benefits greatly from measuring over each point, or voxel column, of the leaf area, allowing for the integration of millions of thickness measures, thus buffering local errors due to false segmentation. Volumetric anatomical traits became constant in variation and values at a minimum of 5 training slices for the bundle sheath, mesophyll cells, and the airspace. As with precision and recall, vein volume substantially varied until about 7 training slices, where values and variation plateaued. The volume of the leaf, the whole mesophyll and epidermises each exhibited similar results as for their thickness.

**Figure 3.**
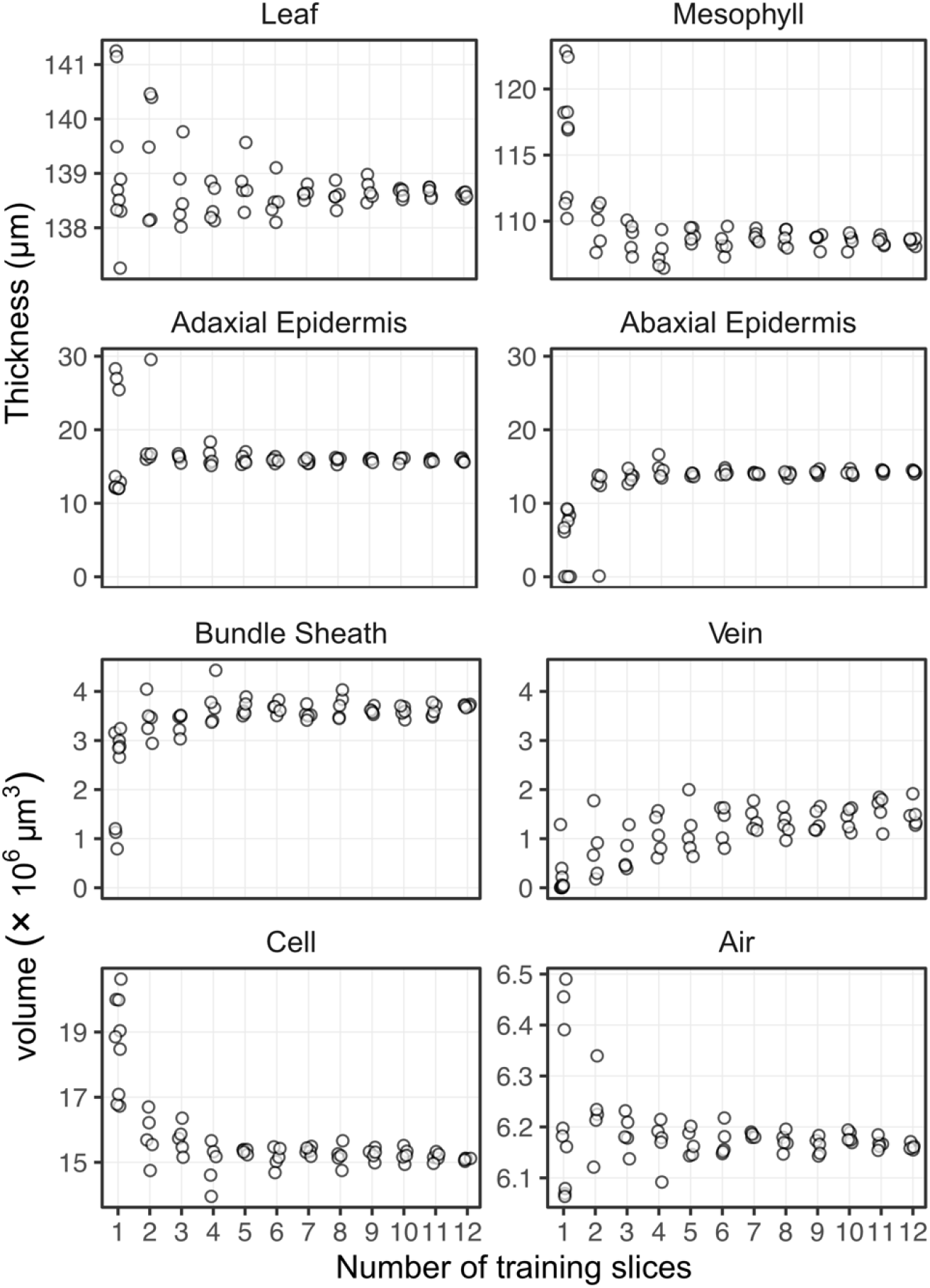
Variation in the measured tissue thicknesses and volumes based on the number of training slices used. False classification of inner leaf pixels as epidermis occurs more with one or two training slices, which resulted in the two epidermis being connected together, hence making the individual thickness estimates wrong. Standard deviations of the thickness estimates are presented in Supplemental Figure S1.

### How many slices should be hand labeled?

In the test presented above, the greater the number of total pixels represented by any class, the fewer training slices required to reach maximum sensitivity (i.e. recall). For example, the air, cells, and background classes are the most common pixel types and clearly show >90% sensitivity (recall) training on as few as 2 manually segmented slices (Figure 2), and with 4 slices different models generated similar biological traits (Figure 3). Veins and bundle sheaths are difficult to segment as they generally present very low contrast between each other and are difficult to distinguish both computationally and visually. In the usage we have made of the program, we are generally interested in defining where the vasculature (veins and bundle sheath together) is rather than extracting traits related to those tissues, and as such 6 slices seem appropriate to get a reliable prediction of the volume of those two tissues (Figures 2 and 3). More problematic is when a class is represented by a smaller number of pixels, the number of training slices required to reach maximum sensitivity in this class increases (Figure 2). This is an inherent class imbalance issue that cannot be solved due to the anatomy of the leaves: the number of training slices needed should allow to reach the desired precision and recall for the class with the lowest amount of pixels per slice. For example, thinner tissues like the epidermis require a minimum of 5 training slices to reach constant precision, recall, and biological traits. As the imaged thickness or smaller axis of a tissue is dependent on the magnification used, care should be taken when planning a scanning endeavor to have the right magnification to have enough pixels per tissue or class of interest in order to facilitate subsequent segmentation. Using the number of slices testing above on previous scans could help guide microCT setup.

## CONCLUSION

We present here a image segmentation framework using open source software to automatically segment image stacks of microCT scans consisting of thousands of single images and requiring only a few hand labeled single slices for each scan. This tool has allowed us to considerably speed up the segmentation of leaf scans and with an increased level of detail: a well-trained user can take less than one hour to prepare the model training slices needed to segment a whole 3D scan and extract relevant biological information. As a comparison, the coarse hand segmentation done in Théroux-Rancourt et al. (2017) took about one full day of work for a pixel volume about a quarter of the size we are presenting here. Further, this segmentation and analysis pipeline has been successfully used on a variety of species and leaf forms (e.g. deciduous and evergreen laminar leaves, C3 grass leaves, conifer needles; Théroux-Rancourt et al., 2020), and is not limited to the tissues extracted above (e.g. resin canals in Figure 4). While our objective was not to provide a universal tool to segment with a single model multiple leaf types, species, and scanning sessions, we consider that this framework has the ability and promise to be adapted and used on other plant material, such as different types of seeds, fruits, stems, and roots, to produce high quality segmentations.

**Figure 4.**
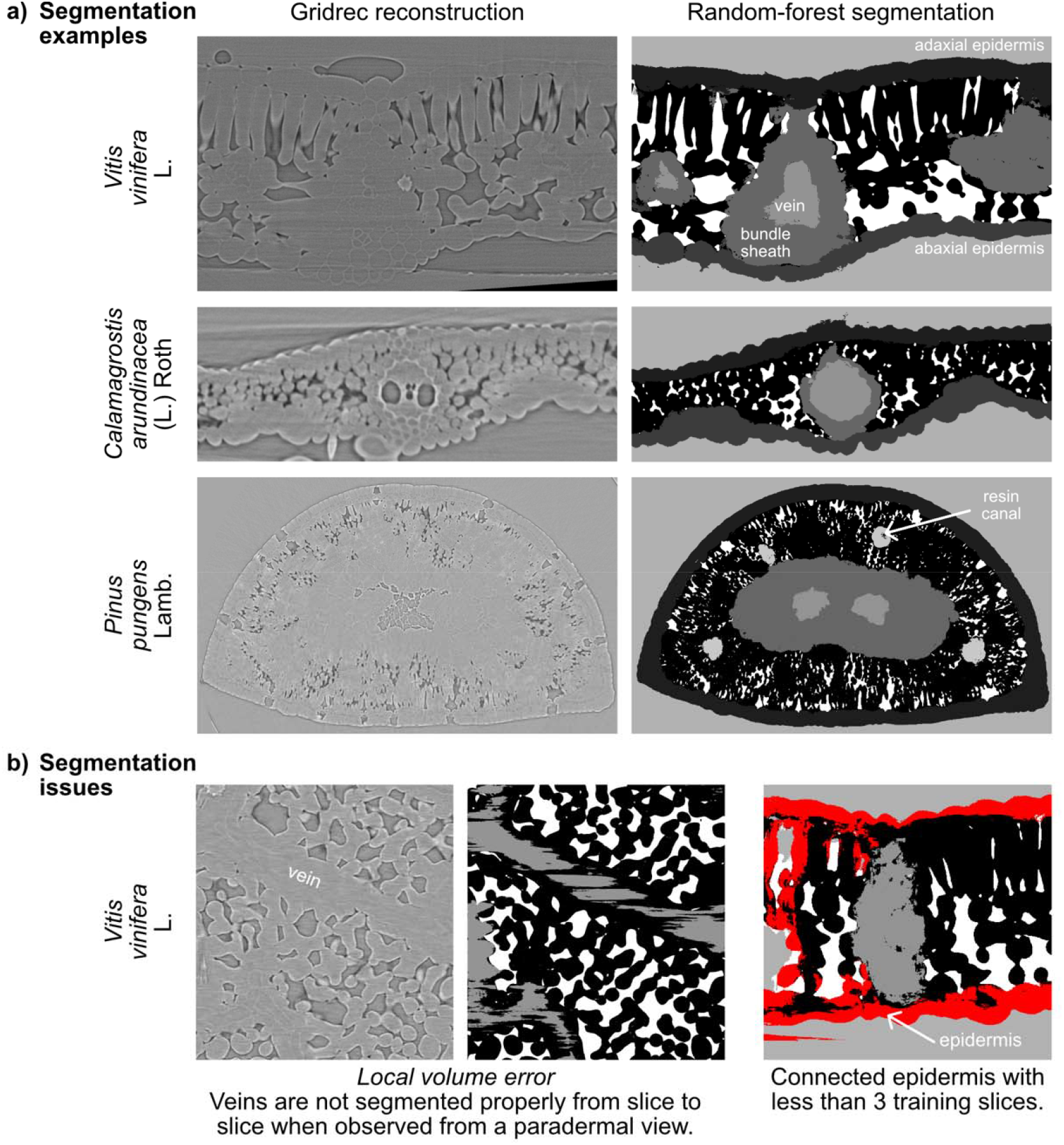
Random-forest segmentation examples (a) for a grapevine leaf, a grass, and a pine needle, and common segmentation issues (b). Gridrec reconstructions on the same slices are shown on the left to compare with the predicted tissues based on random-forest models trained on hand-labelled slices. One of the main segmentation issues is the local volume problem, caused by 2D rather than 3D segmentation, where for example veins are labelled on one slice and not on the other (black areas in between grey-labelled veins). Another issue is having the epidermis connected throughout the leaf at small number of model training slices, here highlighted in red, where a volume might appear disconnected in 2D but is connected in 3D.

However, this framework currently has limitations. For example, certain tissues are not evenly segmented, do not present the expected biological pattern, or present local volume errors, such as veins and bundle sheath constricting and expanding where they should be even from slice to slice. Using a slice-by-slice, 2D model training and segmentation approach can enhance this, and other machine learning methods can probably perform better on this front (e.g. Çiçek et al., 2016). However, we provide a simple tool that can be run on a regular workstation, without the requirement of special infrastructure such as a GPU cluster, for example. This tradeoff was acceptable for the majority of our work. Further, models are currently generated for single scans and have yielded poor results when applied to other scans even of the same scanning sessions and the same species (i.e. similar settings and material). Again, this was an acceptable tradeoff as it significantly sped up the processing of microCT scans as mentioned above. Finally, while it requires more intensive annotation or hand labelling than the common routine for other machine learning tools such as Ilastik (Berg et al., 2019) and Trainable Weka Segmentation (Arganda-Carreras et al., 2017), such tools give poor results on our images when sparsely annotated, most probably because of the limited or absence of gray value contrast between biologically-relevant tissues. Future milestone would be to implement 3D learning to better account for continuous and regular tissues, make the trained model usable for similar scans (e.g. same sessions, species, and material), and testing the performance of other classifiers like support vector machine, k-nearest neighbors, and naive bayes

To conclude, this segmentation framework allowed us to generate a considerable amount of segmented leaves over a wide array of species (see Théroux-Rancourt et al. 2020), and empowers researchers to broaden sampling, to ask new questions about the 3D structure of leaves, and derive new and meaningful metrics for biological structures.

## ACKNOWLEDGEMENTS

We would like to thank Klara Voggeneder for hand labelling the microCT test scan and for improving the hand labelling method, Mina Momayyezi for testing the program, Santiago Trueba for testing the program and providing the images for *Pinus pungens*, and Goran Lovric at the Swiss Light Source for assistance during beamtime. GTR was supported by the Austrian Science Fund (FWF), project M2245.

## AUTHORS CONTRIBUTIONS

JME and MRJ conceptualized and programmed the random forest segmentation program, with assistance from GTR. GTR programmed the leaf traits analysis and the segmentation testing, with assistance from MRJ. GTR acquired and processed the microCT test scan at the SLS. EJF tested and commented on the segmentation program during the development. GTR, MRJ, and JME wrote the paper, with revisions from all authors. AM and CRB provided funding and access to data.

## DATA ACCESSIBILITY STATEMENT

The code and an in depth user manual is available online at github.com/plant-microct-tools/leaf-traits-microct. Future updates will be integrated to this repository.

