## Supplementary figures and images for "Digitally Deconstructing Leaves in 3D Using X-ray Microcomputed Tomography and Machine Learning"

### Supplemental Figure S1

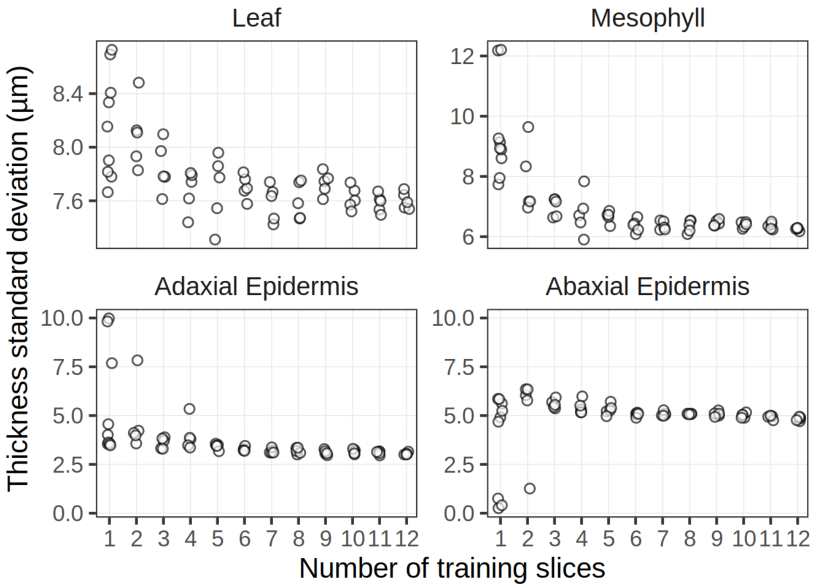


**Figure S1.** Standard deviation of thickness estimates presented in Figure 2.
