## Supplementary Table S1 for "Digitally Deconstructing Leaves in 3D Using X-ray Microcomputed Tomography and Machine Learning"

**Supplementary Table S1.** Average proportion of pixels per tissue in the 24 slices of the training dataset.

| | Number of items[a] | Proportion of pixels in slice | |
|---|---|---|---|
| | | Average | Min-Max |
| Background | 2 | 0.295 | 0.201-0.325 |
| Airspace | - | 0.131 | 0.089-0.182 |
| Mesophyll cells | - | 0.321 | 0.270-0.417 |
| Epidermis | 2 | 0.159 | 0.144-0.180 |
| Bundle sheaths | 1-4 | 0.082 | 0.051-0.132 |
| Veins | 1-4 | 0.014 | 0.005-0.042 |

[a]: Number of separate items sharing the same label present in the slice. For airspace and mesophyll cells, generally all the pixels are connected, with many small regions disconnected.
